# ESMAdam: a plug-and-play all-purpose protein ensemble generator

**DOI:** 10.1101/2025.01.19.633818

**Authors:** Zongxin Yu, Yikai Liu, Guang Lin, Wen Jiang, Ming Chen

**Affiliations:** Department of Engineering Sciences and Applied Math, Northwestern University, Evanston, IL, 60201; Department of Mechanical Engineering, Purdue University, West Lafayette, IN, 47907; Department of Chemistry, Purdue University, West Lafayette, IN, 47907; Department of Biochemistry and Molecular Biology, The Pennsylvania State University, University Park, PA 16802 USA

## Abstract

Proteins often adopt multiple ensemble conformations to perform essential functions such as catalysis, transport, and signal transduction. Traditional physics-based methods for generating these conformations, including molecular dynamics and Monte Carlo simulations, are computationally expensive and time-consuming, limiting their practicality for high-throughput applications like screening. Recent advances in machine learning, particularly deep generative models, offer a promising alternative for protein conformation ensemble generation. However, these models are often task-specific or rely on strong assumptions to generalize. Here, we introduce ESMAdam, a versatile and efficient framework for protein conformation ensemble generation. Using the ESMFold protein language model ESMFold and ADAM stochastic optimization in the continuous protein embedding space, ESMAdam addresses a wide range of ensemble generation tasks. In this work, we demonstrate several basic applications of ESMAdam, including conditional ensemble generation and CG-to-all-atom backmapping. In addition, we showcase advanced applications, such as screening alternative binding modes of protein multimers and reconstructing 3D structures from cryo-EM images. Compared to traditional physics-based methods, ESMAdam significantly reduces computational time. Unlike deep-generative-model-based approaches, it requires no retraining and easily adapts to diverse ensemble restraint conditions, making it exceptionally suited for various structure prediction and screening tasks. This plug-and-play framework represents a step toward efficient and flexible protein ensemble generation for applications in structural biology and drug discovery.

## Introduction

The inherent dynamic nature of proteins is crucial for modulating their activity, facilitating interactions with other biomolecules, and regulating intricate biological processes such as signal transduction and molecular transport. Generating and analyzing protein conformational ensembles is essential for capturing this dynamism, as single static structures fail to represent the full spectrum of functional states. Accurate ensemble generation provides critical insights into the mechanisms underlying protein behavior in various states, enabling advancements in structural biology, drug discovery, and protein design.

Protein conformation ensemble generation consists of a diverse tasks, each tailored to address specific scientific or practical needs. One fundamental objective is to generate protein conformational ensembles that follows the Boltzmann distribution, reflecting the thermodynamic stability of the system under physiological conditions.^1–4^ Beyond thermodynamic considerations, many tasks focus on generating conformation ensembles that satisfy specific geometric or functional constraints. For instance, these may involve generating conformations that facilitate interactions with ligands^5,6^ or protein clustering, ^7,8^ stabilize particular secondary structures, ^9^ or adopt specific topologies critical for biological function. ^10^ Another critical task is protein conformation inpainting, which involves reconstructing missing sections of a protein structure by simultaneously conditioning on its sequence and the surrounding structural context. This task is particularly relevant in coarse-grained molecular modeling, where atomistic details are often simplified for computational efficiency, leading to a gap between coarse-grained structures and atomistic-detail requirements in downstream applications, such as force field refinement, functional analysis, and experimental validation. Acurately recovering missing atomistic details, named as a “backmapping” process,^11–13^ ensures the integrity of the resulting structures and enhances their utility for downstream applications. Finally, generating protein conformation ensembles from cryo-electron microscopy (cryo-EM) images represents another significant challenge.^14–17^ Cryo-EM records proteins and complexes in various functional states. However, interpreting cryo-EM data often focuses on resolving the most stable structure and recovering the underlying conformational ensemble remains complex. Accurate ensemble generation from cryo-EM images involves mapping 2D electron density projections of individual molecules to 3D structures while accounting for experimental noise and heterogeneity. This capability is transformative for understanding flexible and multi-state proteins, as it enables the reconstruction of complete conformational landscapes.

Advancements in AI-driven methods, including AlphaFold, ^18,19^ RoseTTAFold, ^20^ trRosetta, ^21^ and ESMFold,^22^ have significantly improved the accuracy and efficiency of protein structure prediction. More recently, the focus has shifted from single-structure predictions to generating protein conformational ensembles and capturing the dynamic range of protein states. Early approaches, such as MSA subsampling^23^ and clustering, ^24^ expanded AlphaFold’s output by modifying model inputs to produce a more diverse set of conformations. Advanced methods have used deep generative models, particularly diffusion models, ^25,26^ to address the challenge of ensemble generation. ^27–30^ These models utilize stochastic perturbation processes to effectively explore the conformational landscape, enhancing both accuracy and diversity, while predicting the conformational dynamics that govern protein behavior. Deep generative models have also been broadly applied to other conformation generation tasks, including protein structure inpainting,^31–35^ protein-ligand docking prediction, ^36–38^ and protein-protein interaction modeling.^39^ However, these models are typically designed for one or a few specific conformation generation tasks. Recent advances have demonstrated that diffusion models can be adapted for general-purpose ensemble conformation generation tasks with arbitrarily defined ensemble constraints. ^40^ Despite this versatility, the performance of such models declines significantly as the nonlinearity and dimensionality of the ensemble constraints increase, limiting their ability to generalize across complex tasks.^41^

In this study, we introduce ESMAdam, a simple yet efficient method for general-purpose protein conformation ensemble generation. Building upon the pretrained protein language model ESMFold,^22^ ESMAdam utilizes Adam stochastic optimization^42^ on the high-dimensional embedding space of protein sequences. This approach is based on the assumption that protein conformation ensembles are latently embedded near the native structures within the high-dimensional embedding space. Unlike other single-purpose protein conformation generation models, ESMAdam is highly flexible and can accommodate a wide range of tasks. We demonstrate its efficacy through extensive protein conformation generation experiments, focusing on four key applications: (1) controllable conditional conformation ensemble generation for a variety of stable, fast-folding, and intrinsically disordered proteins, (2) CG-to-all-atom configuration backmapping, (3) screening alternative binding modes of flexible protein-protein interactions, and (4) reconstructing protein 3D structures from cryo-EM images. Numerical evaluations reveal that ESMAdam consistently achieves high performance. We propose that ESMAdam serves as a versatile computational tool for rapidly generating protein ensembles, with broad applications in structural biology and drug discovery.

## Results

### Overview of the ESMAdam model

Fig. 1 summarizes a high-level methodology framework of ESMAdam. The core philosophy of ESMAdam is that protein conformation ensembles can be encoded in the latent space near the embedding of the native structure. if a protein conformational ensemble is characterized by low-dimensional features such as experimental ensemble measurement, stable protein structures constrained by low-dimensional features are encoded in the latent space. By exploring the latent space variable with constraints from low-dimensional features, it is possible to generate reasonable protein conformation ensembles. With the recently developed protein language model ESMFold, which excels in predicting native protein structures, ESMAdam leverages the latent space as a trainable variable. For any given ensemble constraint, ESMAdam iteratively updates this trainable latent space variable using stochastic gradient descent methods, such as Adam, ensuring the generated protein conformations align with the desired constraints. Despite its simplicity, this method is highly adaptable to a wide range of tasks.

**Figure 1:**
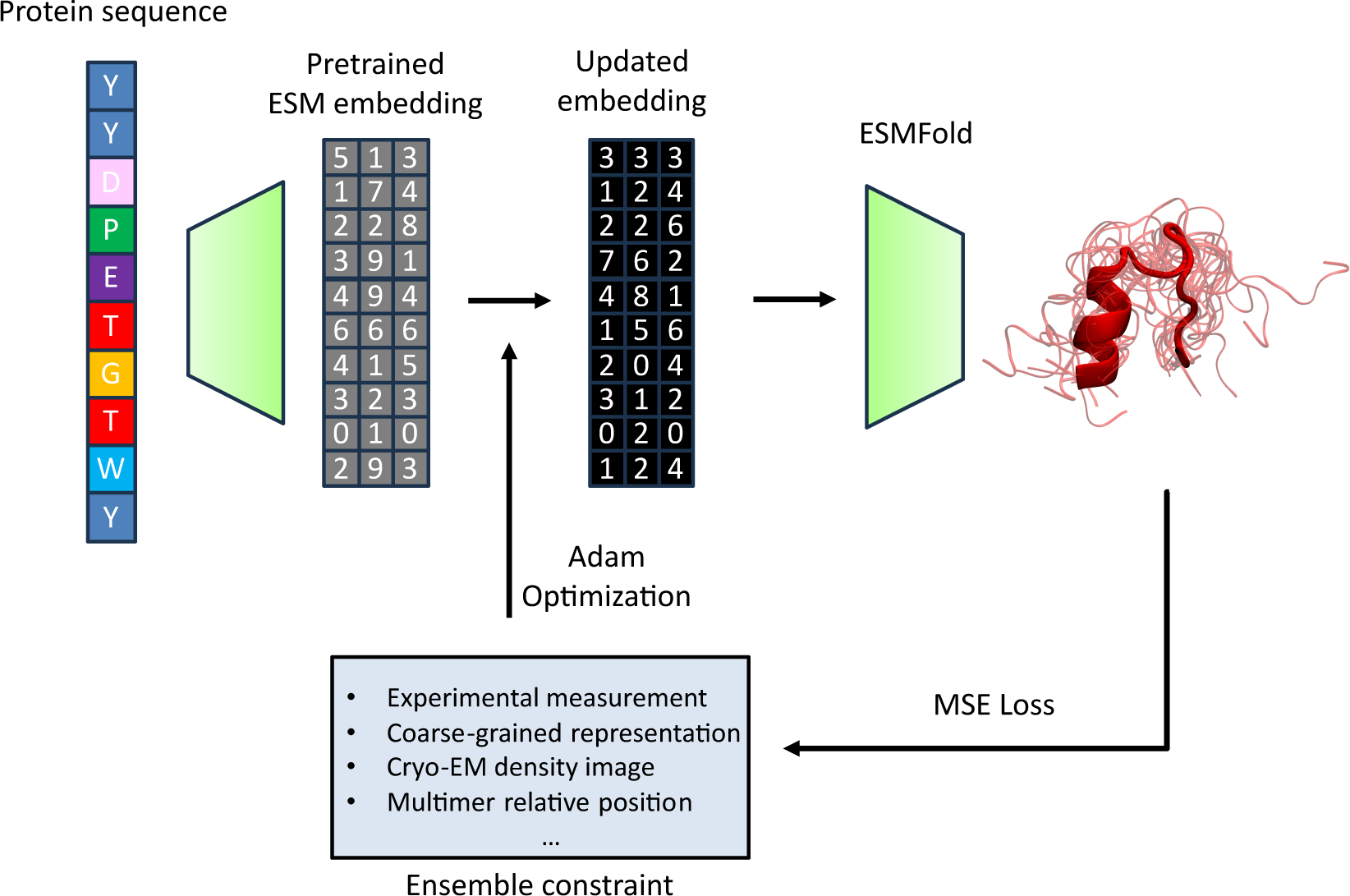
The high level framework of ESMAdam. In ESMAdam, the protein sequence of interest is first embedded using the pretrained protein language model ESMFold. This embedding represents the value in the latent space corresponding to the native structure of the sequence. ESMAdam treats the embedding space as a trainable parameter. The embedding is passed through the trunk of ESMFold to generate the corresponding 3D protein structure. To generate conformation ensembles under specific constraints, the embedding parameter is optimized using a Mean Square Error (MSE) loss function. Optimization is performed with a stochastic gradient descent method, such as Adam. The embedding parameter is updated iteratively until the loss converges below a defined threshold, resulting in a physically plausible ensemble of protein conformations that align with the desired constraints.

### Conditional protein conformation generation

We demonstrate the effectiveness of ESMAdam on conditional protein conformation ensemble generation on a comprehensive benchmark dataset, which consists of a diverse set of proteins, including ordered (BPTI, gb3, Ubq), fast-folding (BBA, BBL, Homeodomain, ProteinB, TrpCage and WWDomain), and intrinsically disordered proteins (PaaA2, drkN, RS-peptide). This dataset spans a wide range of structural characteristics, including varying degrees of order, secondary structure compositions, and sequence lengths. Protein ensembles representing the ground truth Boltzmann distribution were obtained from long, unbiased molecular dynamics (MD) simulations. Previous studies^40^ have shown that diffusion-model-based protein ensemble generation models can generate reasonable protein conformation ensembles when guided by both global and local feature distributions. Following this frame-work, we generated protein conformation ensembles guided by two key features: radius of gyration and secondary structure. These features are experimentally accessible through techniques such as small-angle X-ray scattering (SAXS),^43–47^ nuclear magnetic resonance (NMR), ^48–50^ and circular dichroism spectroscopy^51–55^ experiments. For fair comparison, in this experiment, the distributions of these features are directly obtained from the reference MD simulation. To evaluate the generated ensembles quantitatively, we compare the equilibrium free energy surfaces between ESMAdam-generated conformation ensembles and ground-truth MD simulations. These free energy surfaces were generated by projecting protein configurations to UMAP collective variables derived from ground truth MD simulations. The result, as shown in Fig. 2, highlights the ability of ESMAdam to accurately capture the conformational diversity and thermodynamic properties of the protein ensembles.

**Figure 2:**
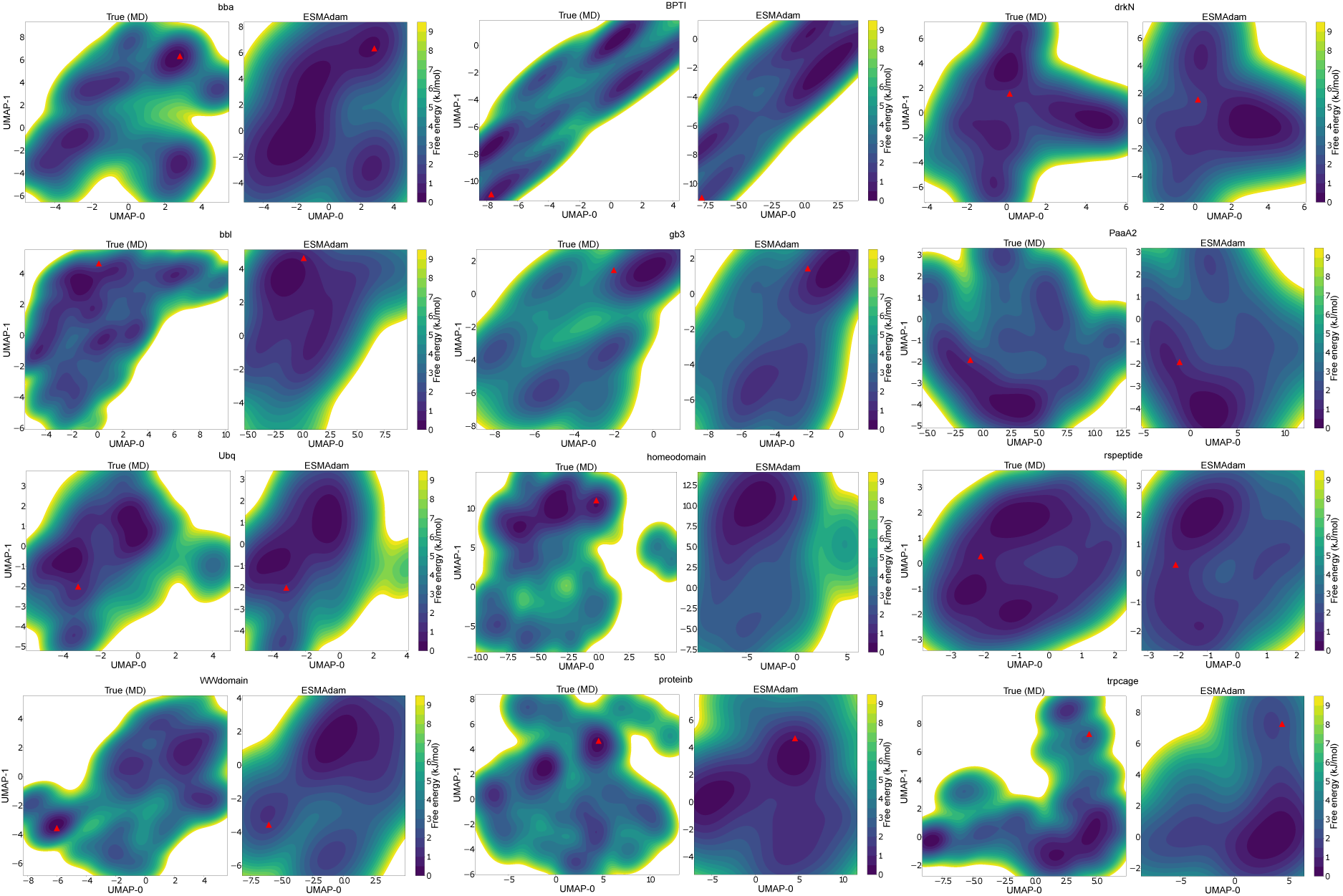
Comparison between the reference MD simulation (left) and ESMAdam (right) guided by radius of gyration distribution and secondary structure free energy surface across the two dimensional UMAP for each protein. The UMAP mapping function was parameterized with the backbone torsion angles of the conformation ensembles from the reference MD simulation. The red triangle represents the native structure predicted by the ESMFold.

### CG-to-all-atom configuration backmapping

Coarse-grained (CG) models are important for studying protein structures, thermodynamic properties, and conformation dynamics. However, the coarse-graining process inherently results in the loss of detailed atomic information, making protein backmapping, which reconstructs an all-atom ensemble from a CG configuration, a critical step in downstream applications. Recent advances in data-driven backmapping, particularly with generative models, have facilitated efficient methods that bypass computationally expensive physical simulations. Despite these advancements, the diversity of CG models,^56–62^ ranging from high-resolution multi-bead-per-residue representations^58^ to low-resolution ultra-coarse-grained (ultraCG) models^63^ where a single bead represents multiple residues, presents a significant challenge for backmapping. No universal backmapping approach currently combines high accuracy with adaptability across different CG resolutions. In this work, we demonstrate that ESMAdam addresses this gap by adapting to CG models of varying scales without retraining. We conducted two CG-to-all-atom backmapping experiments to showcase its versatility. In the first experiment, each residue was represented by its *α*-carbon position. In the second experiment, an ultraCG representation was used, with a single bead representing five residues. Both experiments were performed on a test set comprising proteins BBA, BBL, Homeodomain, ProteinB, TrpCage and WWDomain, which exhibit multiple states, diverse secondary structures and various sidechain structures. Similar to the last example, we also use MD simulations as the ground truth and to construct the marginal distribution of CG beads. The results of the first experiment are shown in Fig. 3, where ESMAdam successfully recovers sidechain distributions with high fidelity, capturing most modes of sidechain torsion angles. The results of the second experiment, presented in Fig. 4, demonstrate that ESMAdam accurately recovers secondary structure conformation ensembles even under ultraCG conditions where most of the backbone information is absent. These findings highlight ESMAdam’s potential as a universal backmapping solution.

**Figure 3:**
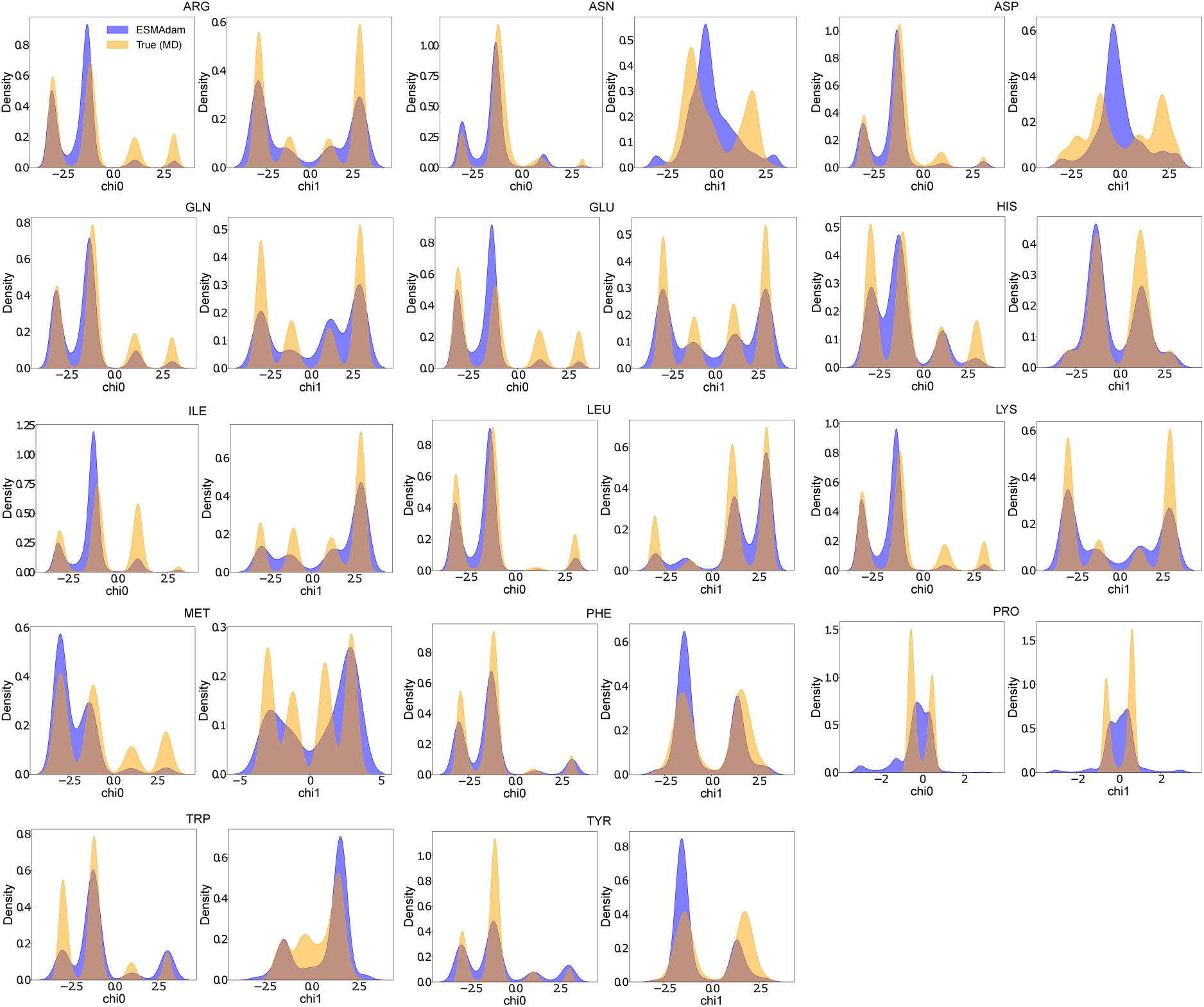
Comparison of the Distribution of Sidechain Torsion Angles: comparison of the distributions of the first (chi0) and second (chi1) sidechain torsion angles for each amino acid type between reference molecular dynamics (MD) simulations and all-atom conformation ensembles generated by ESMAdam from residue-level coarse-grained representations (each residue is represented by its alpha carbon position). The test set includes proteins BBA, BBL, Homeodomain, ProteinB, TrpCage, and WWDomain.

**Figure 4:**
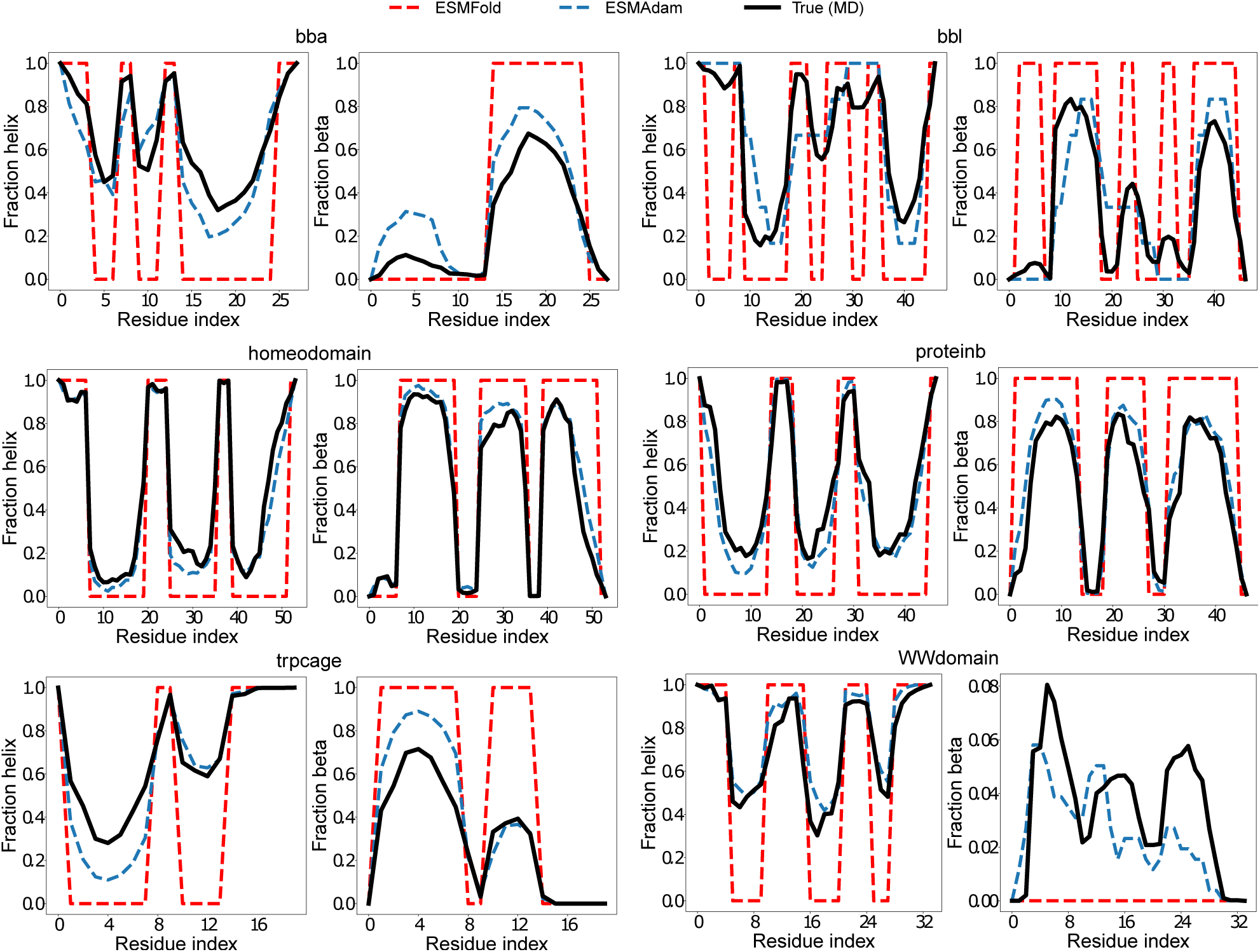
Comparison of the Secondary Structure Percentage Per Residue: comparison between reference molecular dynamics (MD) simulations and all-atom conformation ensembles generated by ESMAdam from ultra coarse-grained representations (where every five consecutive residues are represented as a single bead). The test set includes proteins BBA, BBL, Homeodomain, ProteinB, TrpCage, and WWDomain. To illustrate the secondary structure of the native structure predicted by ESMFold, we also include the secondary structure of the native structure as a comparison.

### Protein complex binding mode exploration

In this experiment, we demonstrate an advanced application of ESMAdam: exploring multiple binding modes in flexible protein complexes. The test system is the well-characterized enzyme–inhibitor complex Barnase-Barstar, a protein complex has been extensively studied experimentally. ^64^ Previous hundreds microseconds molecular dynamics (MD) simulations^65^ have revealed the existence of multiple thermodynamic metastable states approximately 10–20 Åroot-mean-square-displacement (RMSD) away from the native binding structure. Here, we investigate whether ESMAdam can explore and identify both the native binding structure and these metastable states. For this exploration task, the displacement between two proteins’ centers of mass was used as the ensemble constraint, whereas the relative orientation and 3D configuration of each protein are left free to vary. After generating ensembles with ESMAdam, the resulting conformations are relaxed and scored using the Rosetta Suite.^66^ The binding conformation scores from the reference MD simulation and ESMAdam are shown in Fig. 5. The results demonstrate that ESMAdam successfully generates both the native structure and the previously identified metastable states located 20 ÅRMSD from the native structure. Remarkably, ESMAdam also discovers metastable states approximately 5 Å RMSD from the native structure, which were not observed in the elongated MD simulations. These findings highlight ESMAdam’s potential for efficiently exploring multiple binding modes and discovering novel metastable binding states, making it a valuable tool for studying protein-protein interactions and binding energetics.

**Figure 5:**
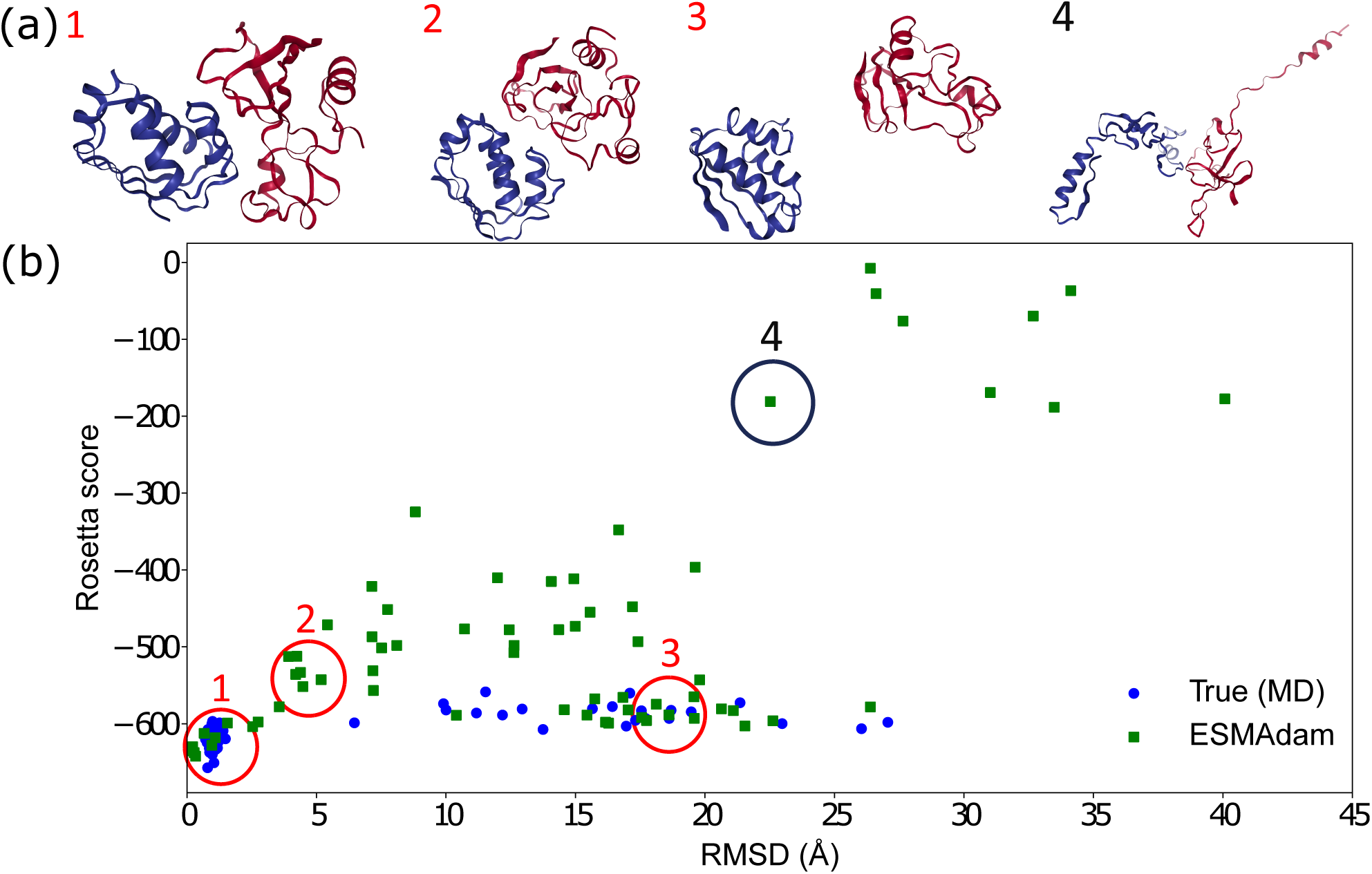
Evaluation of Protein Multimer Ensembles Generated by Reference MD Simulation and ESMAdam: (a) Multiple diverse conformational ensembles generated by ESMAdam, and (b) Rosetta score comparison of ensembles generated by reference MD simulation and ESMAdam. The reference MD simulation uncovers two clusters (1 and 3) of stable states: one corresponding to the native crystal structure and another located 10–20 ÅRMSD from the native structure. In comparison, ESMAdam identifies additional metastable structures (1, 2, and 3) with significantly reduced computational time. Additionally, ESMAdam is capable of generating other less stable structures (e.g., cluster 4), which may provide valuable insights for studying protein multimers under varying biological conditions.

### Reconstructing Heterogeneous Protein 3D Structures from cryo-EM Data

In the final experiments, we extend the capacity of our model to reconstruct heterogeneous protein 3D structures from 2D images obtained by cryo-EM. ^67–72^ Cryo-EM is an important experimental technique for generating high-resolution 3D structures of biological molecules. Recent studies suggested that cryo-EM can potentially extract conformational ensembles following the equilibrium Boltzmann distribution. ^16,73–78^ These methods either use a reconstructed 3D density map as an ensemble constraint for molecular dynamics (MD) simulations, or use converged MD simulations as a prior to conduct ensemble reweighting with 2D density images. However, both approaches are computationally expensive for large proteins. In this experiment, we treat the cryo-EM 2D images as the training objective. Following previous work, ^16^ we adopt a computational model differentiable with respect to the configuration to represent the configuration-to-image formulation process in cryo-EM:

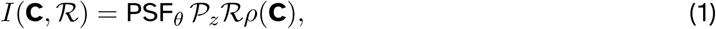

where *ρ*(**C**) is the electron density as a function of the protein configuration **C**, ℛ is the rotation matrix of the electron density, 𝒫*_z_* represents the projection along the *z*-axis, and PSF*_θ_* is a point spread function parameterized by defocus and translation. The electron density *ρ*(**C**) is expressed as the sum of spherically symmetric 3D normal densities centered on each alpha carbon atom of the protein. We applied this computational model to configurations obtained from the reference MD simulation and introduce varying levels of Gaussian noise to generate a synthetic cryo-EM image dataset for each protein. In this experiment, we consider three noise levels: no noise, moderate noise (signal-to-noise ratio of 0.01), and large noise (signal-to-noise ratio of 0.0001). The results, presented in Fig. 6 and supported by additional experiments detailed in the Supplement, highlight several key findings. First, ESMAdam demonstrates superior performance in exploring the protein conformational space, particularly excelling at capturing protein thermodynamic properties. Second, ESMAdam exhibits strong accuracy in predicting secondary structures, as evidenced by its ability to capture the fraction of helical content and beta content. Notably, even at high noise levels, ESMAdam successfully preserves secondary structure predictions and explores the conformational space with high thermodynamic accuracy. Batch optimization inherently mitigates Gaussian noise, which partially explains the high performance of ESMAdam even in the presence of significant noise. These results underscore its robustness to noisy inputs and suggest that ESMAdam holds substantial potential for application to real cryo-EM datasets, which often contain significant noise. However, it is important to emphasize that the noise introduced in this experiment is Gaussian noise with zero mean. For real cryo-EM images, where noise is often non-Gaussian and more complex, the performance of ESMAdam may degrade and might require additional pre-processing or post-processing steps to ensure optimal results.

**Figure 6:**
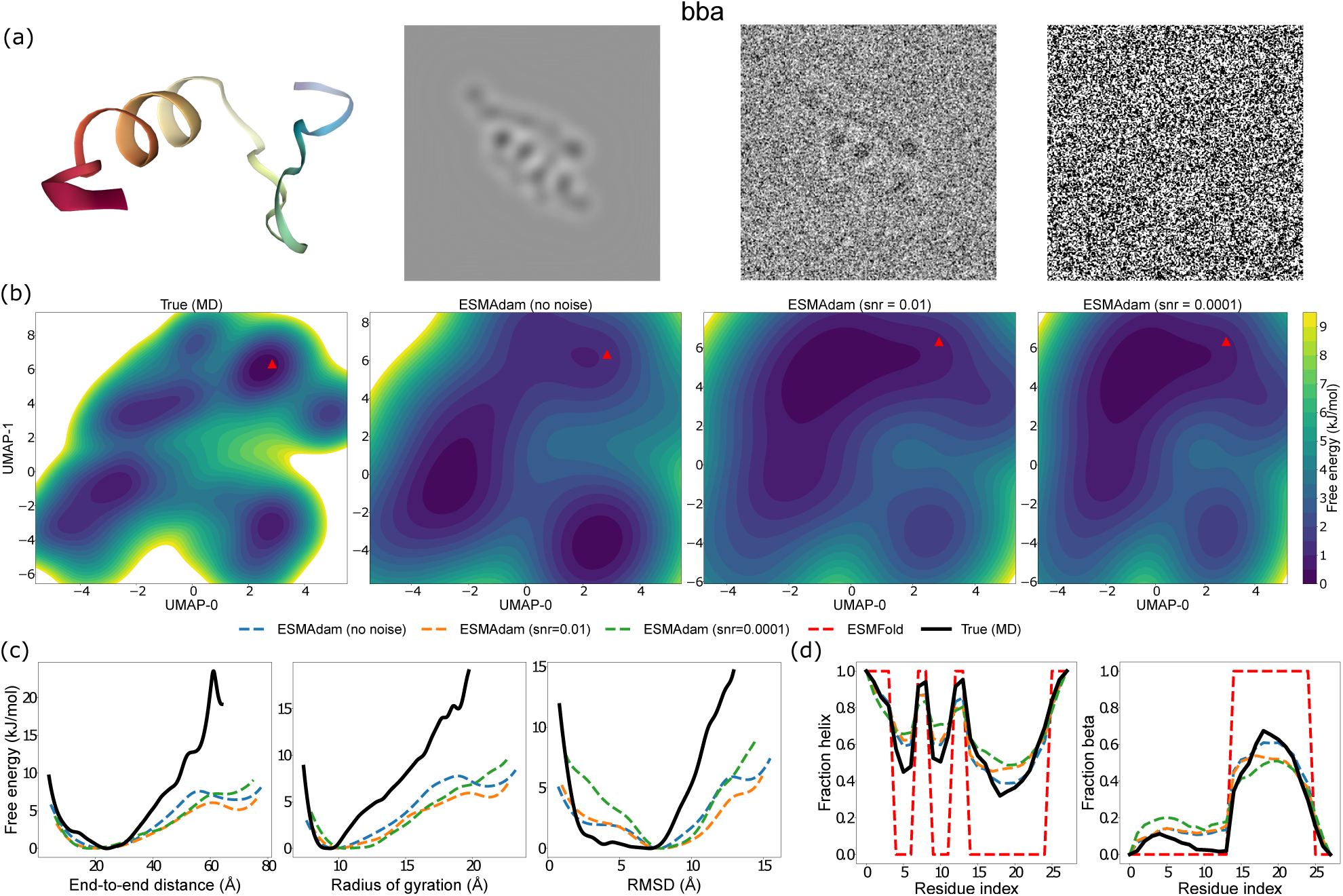
Evaluation of BBA Heterogeneity Reconstructed from cryo-EM Images: (a) Visualization of a BBA structure and its corresponding synthetic cryo-EM images under different signal-to-noise ratios (SNR), from left to right is no noise, SNR=0.01, annd SNR = 0.0001, (b) 2D UMAP free energy surface (FES), (c) physical feature FES, and (d) the secondary structure percentage per residue comparison between reference molecular dynamics (MD) simulations and ESMAdam guided with cryo-EM images under different SNR. Results for other proteins are reported in the supplementary materials.

## Discussion

In this work, we introduce ESMAdam, a general-purpose, plug-and-play method to generate protein conformation ensembles, based on the advanced protein language model ESMFold.

ESMAdam shares conceptual similarities with molecular dynamics (MD) simulations that incorporate ensemble restraints. In MD simulations, ensemble restraints are commonly used to guide conformational sampling, ensuring that the generated ensemble aligns with key features derived from experiments or computational methods. ^79–83^ Although ESMAdam adopts a similar philosophy, it offers a significant advantage by efficiently exploring conformational space. Instead of relying on computationally expensive MD simulations, ESMAdam directly generates diverse protein conformation ensembles guided by ensemble restraints, making it an invaluable tool for protein research, particularly when rapid and accurate ensemble generation is essential.

It is interesting that ESMAdam can generate a protein conformational ensemble qualitatively following the Boltzmann distribution in several examples, although ESMFold is designed to predict a single structure for a protein. Constraints of low-dimensional features define a manifold in the latent space. ESMAdam optimization guides a latent space variable converging to a random point on the manifold, whereas there is no control of the latent space position on the manifold. The success of ESMAdam may be explained by the following reasons. First, most of the latent space around the embedding of the native structure corresponds to “good” structures. Therefore, random sampling in the restraint manifold can still generate thermodynamically metastable structures. Second, the conditional probability distribution of protein structures may be well characterized by a few-mode distribution at room temperature, i.e. only a few low-energy structures dominate the conditional distribution where these structures could be explored by ESMAdam. Verification of these explanations is subject to future studies.

As demonstrated in the Results section, ESMAdam excels at producing physically meaningful protein conformational ensembles under various constraints, showcasing its versatility for downstream tasks. Specifically, we demonstrated its utility in generating high-quality protein conformation ensembles that align with the Boltzmann distribution, performing precise CG-to-all-atom backmapping, exploring multiple binding states of protein complexes, and reconstructing 3D protein ensembles from noisy cryo-EM images. Looking ahead, we anticipate a broad range of additional applications for ESMAdam. For example, by integrating ESMAdam with an energy function, it could be used to identify stable configurations under varying biological conditions and environments, such as protein interactions with materials. ^84^ This versatility underscores the potential of ESMAdam to advance structural biology and expand the horizons of protein research.

Many studies have explored the modification of input to state-of-the-art AI-driven folding models to generate protein ensembles.^23,24^ However, none have achieved the level of quantitative and flexible control or the rigorous thermodynamic evaluation of generated ensembles demonstrated by ESMAdam. Another similar research direction in AI-driven protein conformation generation is the development of fine-tuned models built upon pre-trained foundation protein conformation generation frameworks for specific tasks. For example, recent methods that utilize manifold constraint sampling techniques have been proposed to generate protein conformation ensembles under various ensemble constraints. ^40,78^ Compared to these models, ESMAdam offers several significant advantages. The performance of manifold constraint sampling techniques declines markedly as ensemble constraints become more nonlinear or higher in dimensionality, limiting their applicability to arbitrarily defined or high-dimensional constraints. In contrast, ESMAdam imposes no restrictions on the dimensionality or linearity of the ensemble constraints, provided that they are differentiable with respect to the 3D structure. This flexibility makes ESMAdam suitable for a much broader range of tasks, establishing it as a versatile and robust solution for protein conformation generation.

Despite its superior performance, ESMAdam has a few limitations. A primary concern is that the continuous embedding space to be optimized grows significantly with the size of the protein system, leading to increased GPU memory requirements. This can pose challenges for researchers with limited GPU resources, potentially restricting its accessibility to some users. In the future, our aim is to address this issue by exploring techniques such as Low-Rank Adaptation (LoRA), ^85^ a technique for fine-tuning large-scale models that reduces computational and memory costs by introducing low-rank updates to pre-trained model weights. LoRA has proven to be effective for fine-tuning large language models. By adapting LoRA on the protein embedding space to perform various protein conformation generation tasks, we hope to reduce the memory footprint and computational demands of ESMAdam without compromising its performance. This approach could make ESMAdam more accessible and practical for researchers with varying levels of computational resources, further expanding its utility in the field.

Overall, we have demonstrated that ESMAdam is a powerful, flexible, and easy-to-use tool for generating protein ensembles in a wide range of tasks. We believe that this approach will benefit researchers from diverse fields, including structural biology, computational chemistry, and machine learning. The applications of ESMAdam range from simple tasks such as refining or inpainting protein structures to more complex challenges, such as exploring protein conformational landscapes and high-throughput screening of protein interactions and dynamics. For machine learning researchers, ESMAdam provides insight into the nature of protein metastable ensembles. It reveals that these ensembles may be accessible within the continuous embedding space. This realization opens new avenues for machine learning research, particularly in the development of deep generative models with a focus on embedding-space representations.

## Materials and Methods

### ESMAdam method

A protein language model like ESMFold processes a protein sequence *S*, maps it to a high-dimensional embedding space *E* = embed(*S*), and directly predicts the 3D structure **C** of the native state as *f* (*S, E*) = **C**. ESMAdam extends ESMFold by freezing the pretrained ESMFold parameters and treating the embedding space *E* as a trainable parameter. This approach allows ESMAdam to generate protein conformation ensembles via *f* (*S, E_θ_*) = **C**. Given an ensemble constraint that can be expressed as a continuously differentiable function of the 3D configuration, **r** = *g*(**C**), ESMAdam optimizes the embedding parameters ***θ*** by solving the following objective:

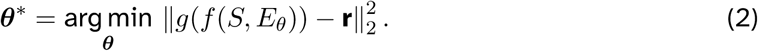

### CG-to-all-atom backmapping experiment

All-atom conformations are represented as **C** ∈ ℝ^3^*^N^*, where *N* is the number of atoms in the protein. We define a linear coarse-graining (CG) mapping function ***ζ*** that reduces the dimensionality of the system: **x** = ***ζ*C** ∈ ℝ^3^*^n^*, where *n* is the number of CG atoms. Various CG mapping methods can be employed. For instance, ***ζ*** can map to the alpha carbon positions or the center of mass of a residue. In this experiment, we consider two types of CG mapping methods. The first maps each residue to its alpha carbon position as the CG representation. The second maps every fifth alpha carbon position (e.g., 0, 5, 10*,…*) as the CG representation. In the CG-to-all-atom backmapping experiment, we define the ensemble constraint as the root mean square distance (RMSD) between the given CG configuration **x** and the CG configuration mapped from the recovered all-atom configuration:

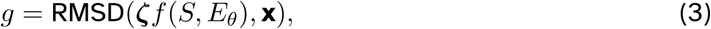

with **r** = 0 in Eq. 2.

### Protein multimer alternative binding mode experiment

The 3D structure of a multimer can be denoted as **C** = {**C**_1_, **C**_2_*, R, t*}, which consists of the 3D structures of each protein monomer, **C**_1_ and **C**_2_, and the relative rotation *R* and translation *t* between the two proteins. In this experiment, the relative translation, represented as the displacement between the centers of mass (COM) of the two proteins, is used as the ensemble constraint, while the remaining degrees of freedom are left free to vary. To ensure that the optimization process is invariant to global rotation and translation, the entire 3D structure is superimposed with respect to a fixed reference structure, **C**_ref_ (e.g., the native structure predicted by ESMFold). The ensemble constraint is then defined as:

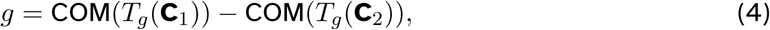

where COM denotes the center of mass operator, and *T_g_* represents the rototranslational operation obtained by superimposing **C** onto **C**_ref_. The ensemble constraint value **r** is a 3-dimensional displacement vector that can either be uniformly sampled from the 3-dimensional space or selected based on prior knowledge to narrow the search space. For simplicity, in this experiment, we assume prior knowledge of the displacement obtained from MD simulations. However, to encourage the exploration of diverse structures, we introduce a small perturbation to the initial embedding value *E*. This perturbation prompts the Adam optimization process to search for alternative 3D structures of the multimer that satisfy the same ensemble constraints, enhancing the diversity of the generated conformations.

### Cryo-EM image as the ensemble constraint

In this experiment, we generate synthetic cryoEM images from snapshots of frames from MD simulation trajectories following the same method from previous works. ^16,40^ The image corresponding to a configuration is generated with its alpha carbon atoms. The COM of the protein is placed at the origin, and a rotation matrix R is placed. The electron density at location **c** = (*c_x_, c_y_, c_z_*) is obtained by summing over 3D Gaussian of width *σ* = 1.5Å centered at each *C_α_* atoms:

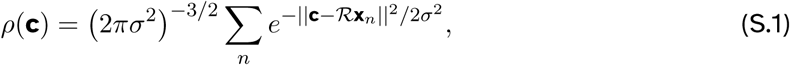

where **x***_n_* is the position of n-th *Cα*. The 2D intensity map *I* is obtained by projecting along the z-axis:

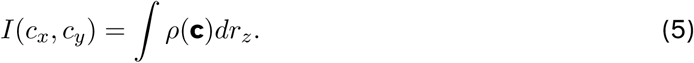

In the experiment, the 2D density map is discretized with 2D grid of size 256×256. The pixel size varies with protein size. In the final step, the 2D density map is convolved with the real-space point-spread function:

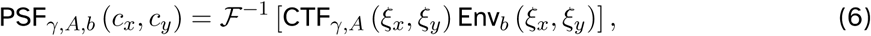

where ℱ^−1^ is the inverse Fourier transform. CTF and Env are contrast transfer and envelop functions:

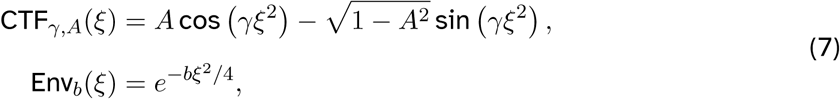

with *ξ_x_* and *ξ_y_* the spatial frequencies, *A* is the amplitude-contrast ratio, *b* is the B-factor, and *γ* = −*π*Δ*zλ_e_* where Δ*z* is the defocus, and *λ_e_* the electron wavelength. In this experiment, we set *A* = 0.1*, b* = 1Å^2^*, λ_e_* = 0.019866Å, Δ*z* uniformly sampled between (0.03, 0.09) *µm*. To construct the dataset of images, we then add Gaussian i.i.d. noise to each pixel with various levels of SNR.

Under the ESMAdam framework, an additional unknown parameter introduced in the 3D reconstruction from cryo-EM 2D density images is the rotation matrix, which we treat as a trainable parameter. Furthermore, we need to ensure that the optimization of the 3D structure remains invariant to global rotation and translation. Thus, given a cryo-EM density image **I**_condition_, we define the ensemble constraint in the experiment as:

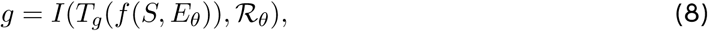

where the image generation function *I* is defined in Eq. 1 and **r** = **I**_condition_. This formulation ensures that the reconstruction process is robust and aligned with the experimental constraints imposed by the cryo-EM image. Our approach shares similarities with ab initio modeling methods in cryo-EM, ^86^ which utilize stochastic gradient descent (SGD) for efficient 3D reconstruction. However, we propose that ESMAdam, leveraging SGD in the continuous embedding space, offers a potential advantage due to ESMFold’s inherent ability to generate stable and accurate structures.

## Acknowledgement

G.L. and Y.L. gratefully acknowledge the support of the National Science Foundation (DMS-2053746, DMS-2134209, ECCS-2328241, and OAC-2311848), and U.S. Department of Energy (DOE) Office of Science Advanced Scientific Computing Research program DE-SC0023161, and DOE – Fusion Energy Science, under grant number: DE-SC0024583.

## Author contributions statement

Y.L. and Z.Y. designed the research; Y.L., and Z.Y. performed research; Y.L. and Z.Y. contributed to the code base; Y.L. and Z.Y. curated the data; G.L. W.J. and M.C. supervision; Y.L., Z.Y., G.L. and M.C. wrote the manuscript.

## Supporting information

### 1 Benchmark Protein Dataset

We assemble a testing dataset of 12 proteins, from order proteins to fast-folding proteins to intrinsically disordered proteins, as shown in Fig. 1. The ground truth Boltzmann distribution was obtained from long unbiased MD simulations. Simulations of BBA, BBL, WW domain, Homeodomain, Protein B, and TrpCage were conducted in previous work using the CHARMM22* force field and TIP3P water model at the temperature of 350KThe simulations were performed using adaptive sampling strategy. Simulations of drkN, PaaA2, BPTI,gb3, and Ubquitin were conducted by DE Shaw research using the a99SB-disp force field for 30 *µs*. The force field parameters were optimized based on experimental measurements to achieve high accuracy in simulation results across various ordered and disordered proteins. The simulation of RS-peptide consists long unbiased simulations of up to 2 milliseconds of simulation time, with 20 unique starting structures, each run with 100 clones of ∼0.9 micro-second each using Amber03ws force fieldand Tip4p/2005 water model. Details of RS-peptide simulation can be referred from previous works. Finally, the simulation of Barnase–barstar protein complex system is conducted by DE Shaw research with Amber ff99SB*-ILDN force field and the TIP3P water model using tempered binding simulations.^65^

**Figure 1:**
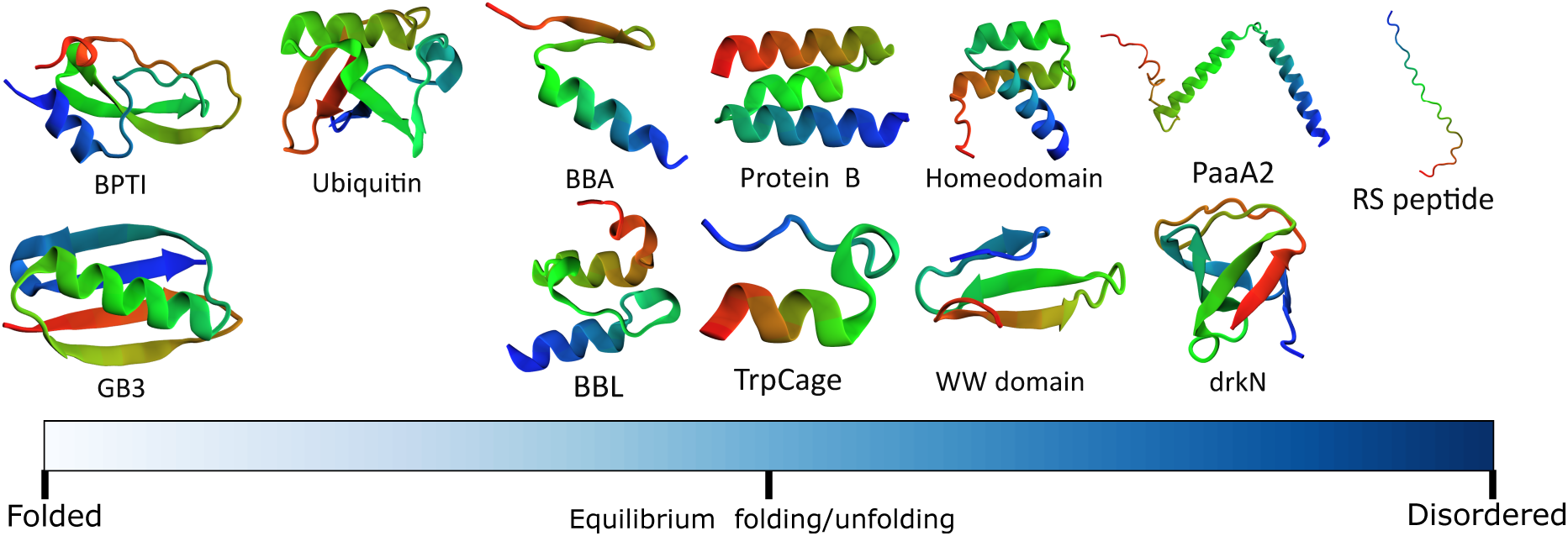
Schematic illustration of benchmark set of proteins used in this work to assess the performance of ESMAdam.

### 2 Full experimental results

#### Condition protein conformation ensemble generation

Here we present the full evaluation of the experiment, including the physical feature FES in Fig. 2 and secondary structure percentage per residue in Fig. 3.

**Figure 2:**
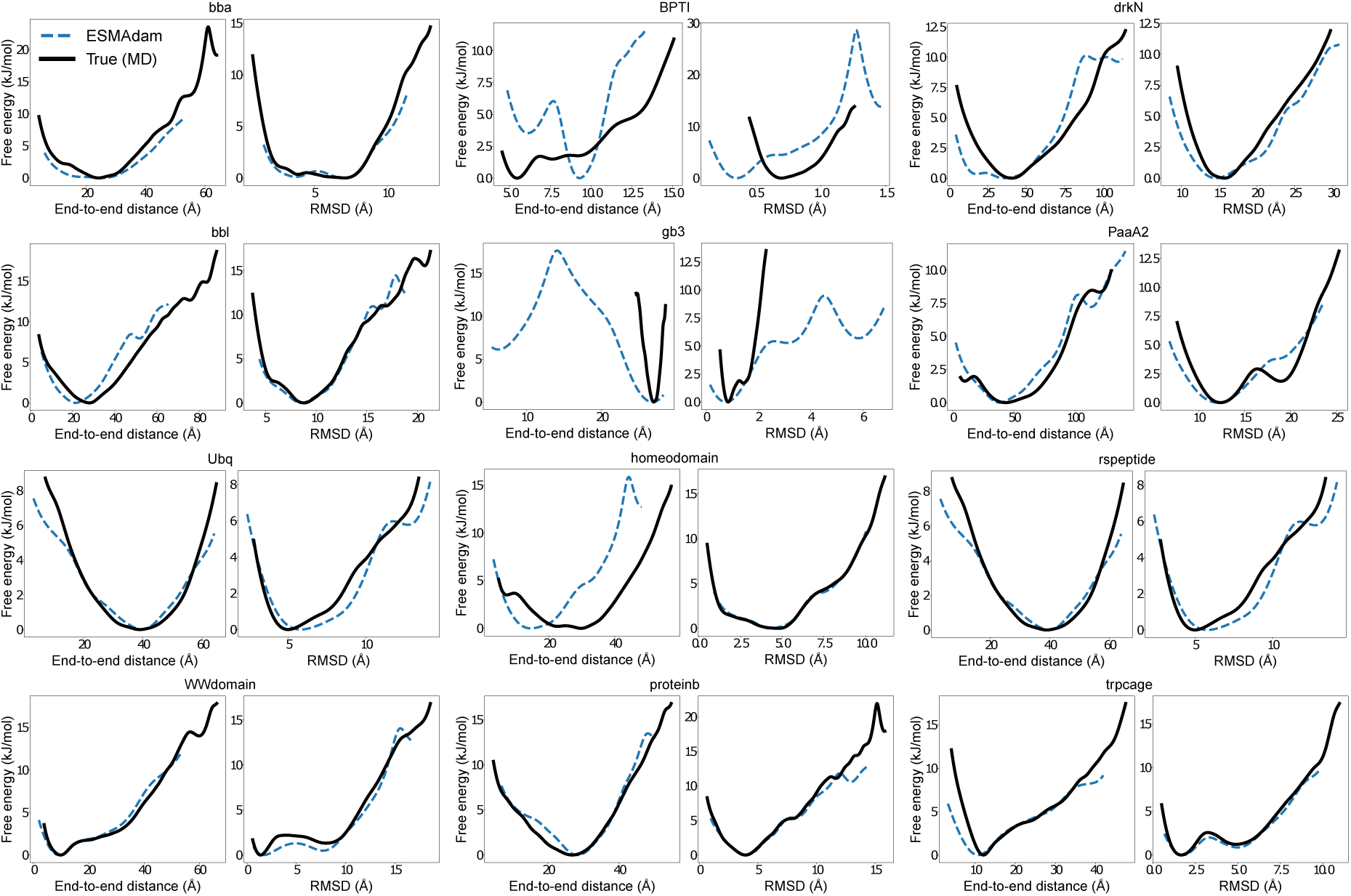
The FES of physical features in all conditional conformation ensemble generation experiments.

#### CG-to-all-atom backmapping experiments

Here we present the full evaluation of the ultraCG backmapping experiment, including the UMAP 2D FES in Fig. 4.

#### 3D structure reconstruction from cryo-EM images

Here we present the additional 3D structure reconstructionn from cryo-EM images experiment for four other proteins: BBL, ProteinB, TrpCage, and WWdomain. The results are shown in Fig. 5, Fig.6, Fig. 7, and Fig. 8.

### 3 Other Methodological Details

#### UMAP parameterization

To analyze protein conformational ensembles, we leverage the backbone torsion angles (*ϕ* and *ψ*) as key structural features. These angles are transformed into a high-dimensional feature space by computing their sine and cosine values to ensure a continuous representation. We represent the protein’s conformational landscape with this high-dimensional representations. To reduce the dimensionality of this space and facilitate visualization, we apply Uniform Manifold Approximation and Projection (UMAP). UMAP is a non-linear dimensionality reduction technique that preserves the local and global structure of the data while projecting it into a lower-dimensional space. By mapping the high-dimensional torsion angle features to a 2D projection, UMAP enables intuitive exploration of the conformational diversity and clustering of protein states. The UMAP function is parameterized with data from the reference MD simulations in all cases.

**Figure 3:**
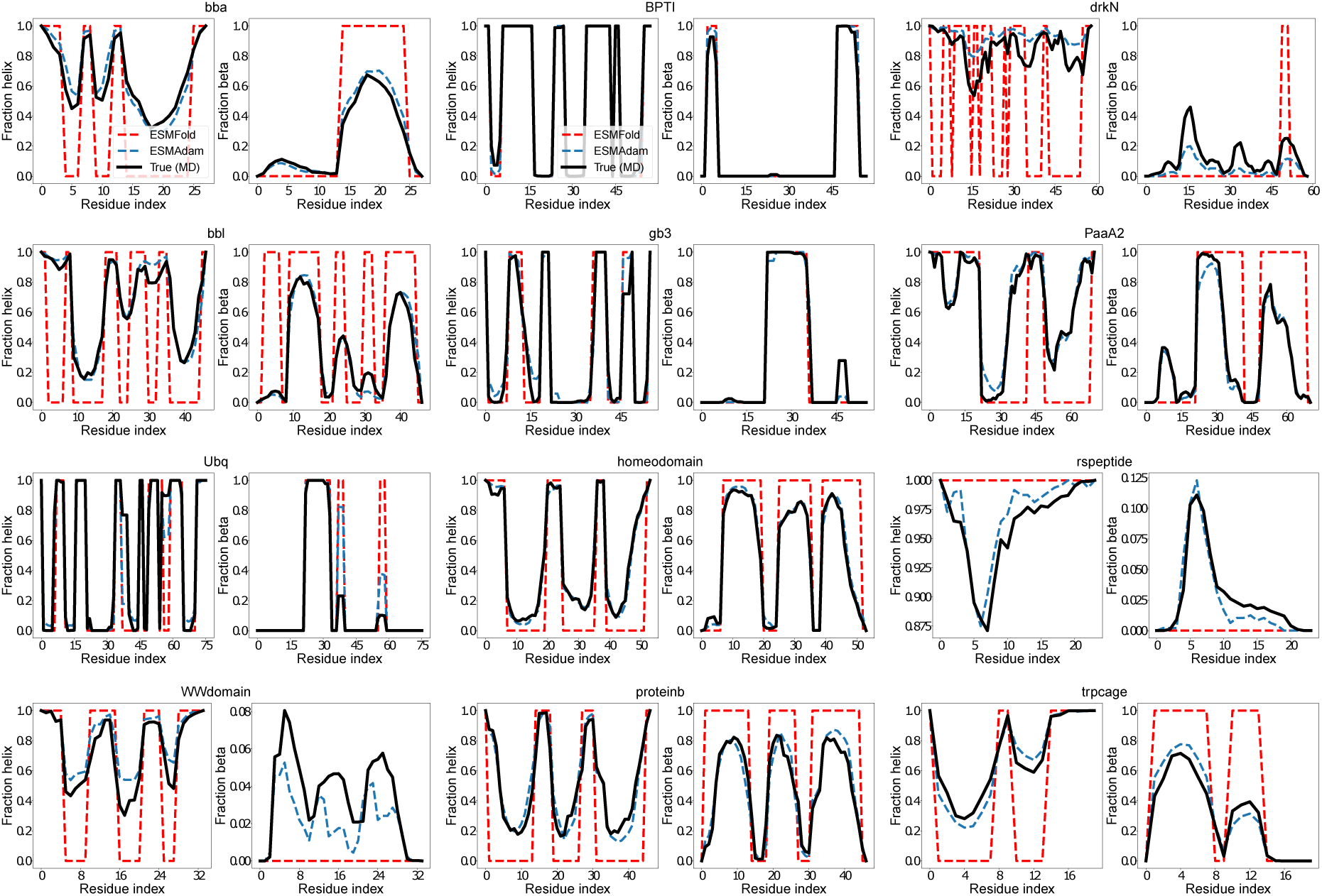
The secondary structure per residue in all conditional conformation ensemble generation experiments.

#### Choice of the correction hyperparameter

An important hyperparameter in ESMAdam is the learning rate. A learning rate that is too low can result in prolonged convergence times, while an excessively high learning rate can cause optimization instability. In this study, we report the empirically determined learning rate values that yield the best performance in Table 1. In most experiments, these empirically chosen learning rates enabled convergence within 100 epochs. However, for the 3D structure reconstruction experiments, convergence required up to 300 epochs, reflecting the increased complexity of this task.

**Figure 4:**
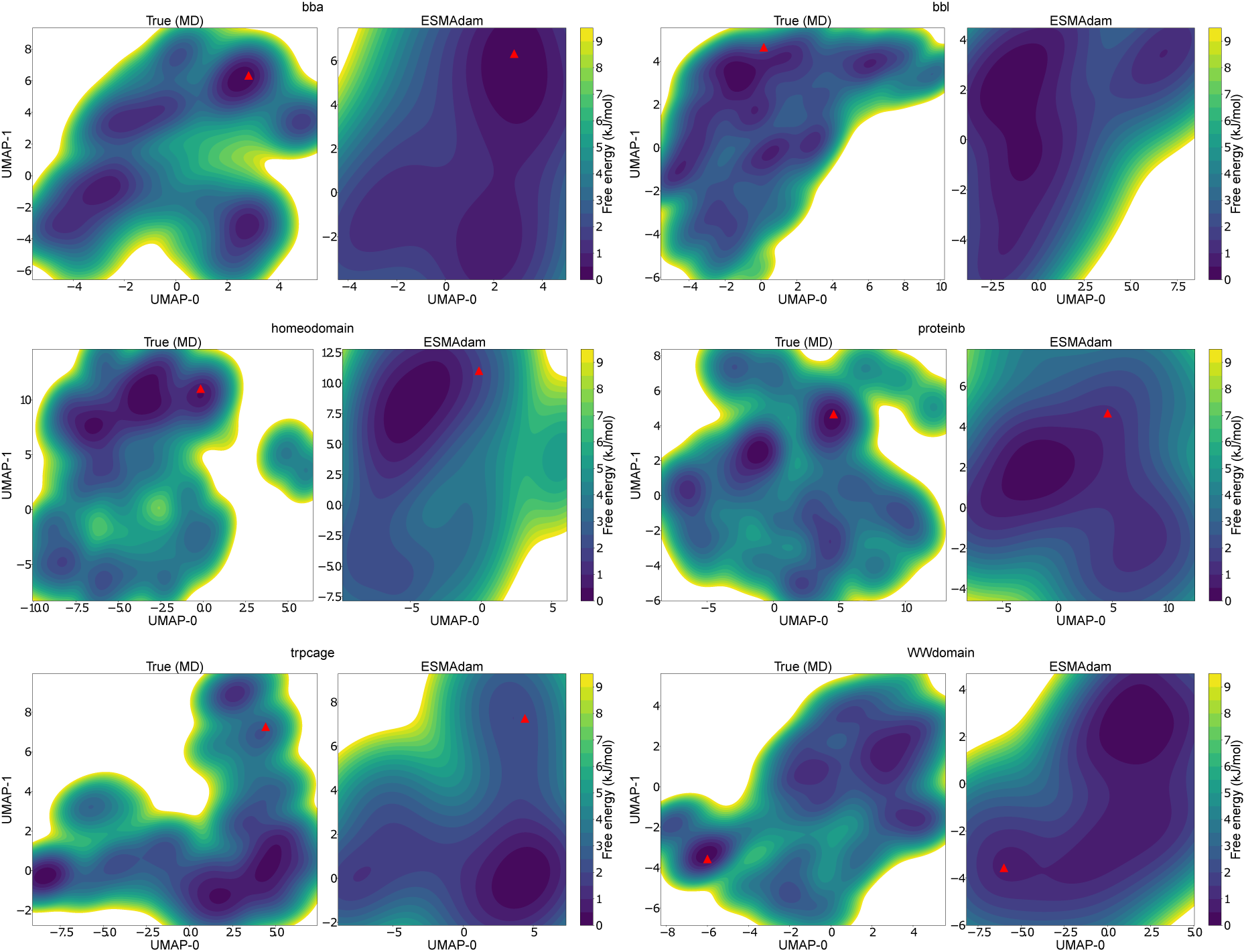
Comparison between the reference MD simulation (left) and ESMAdam (right) in ultraCG backmapping experiment FES across the two dimensional UMAP for each protein.

**Table 1:**
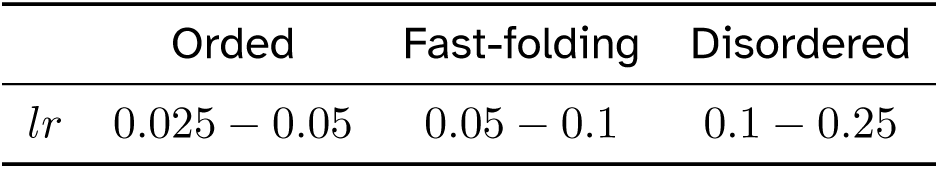
The correction term hyperparameter for each type of proteins.

**Figure 5:**
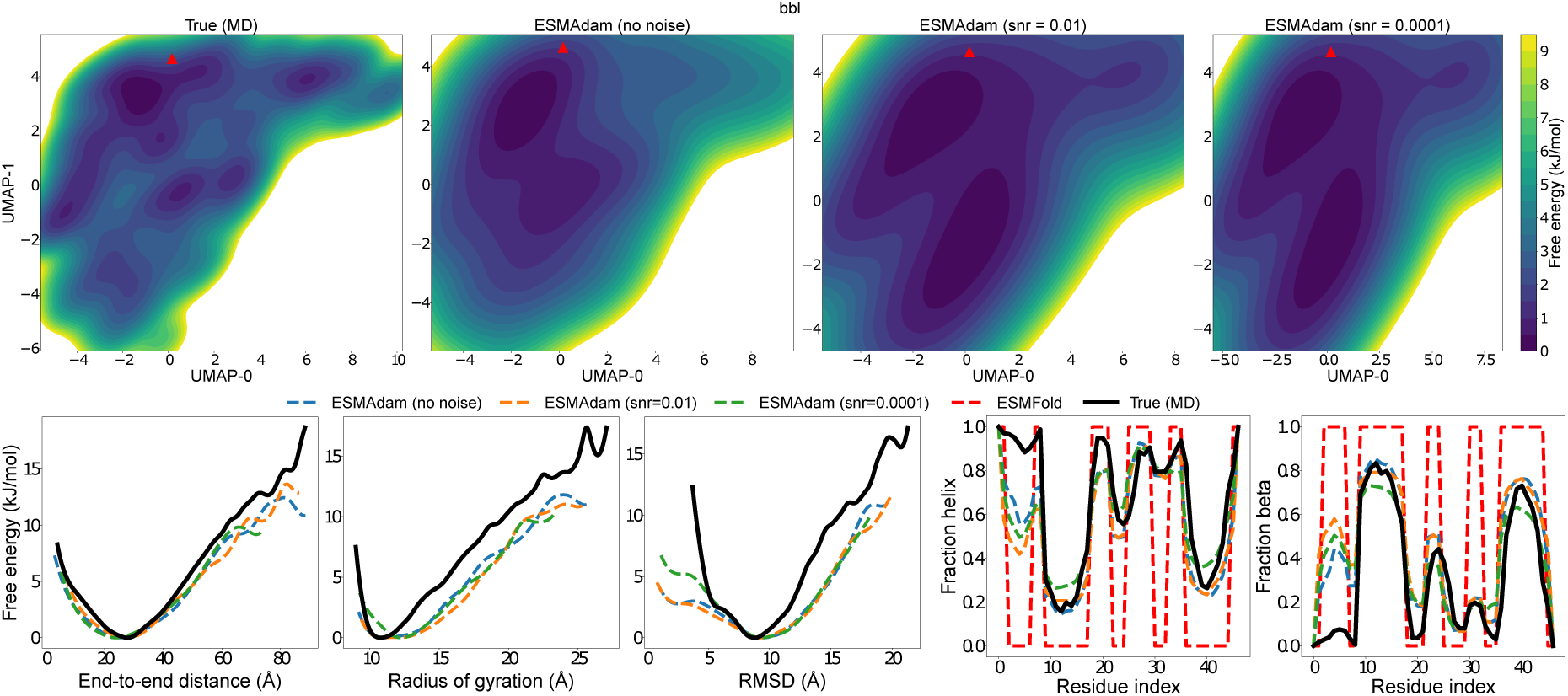
3D conformation ensemble reconstruction from cryo-EM results for BBL

**Figure 6:**
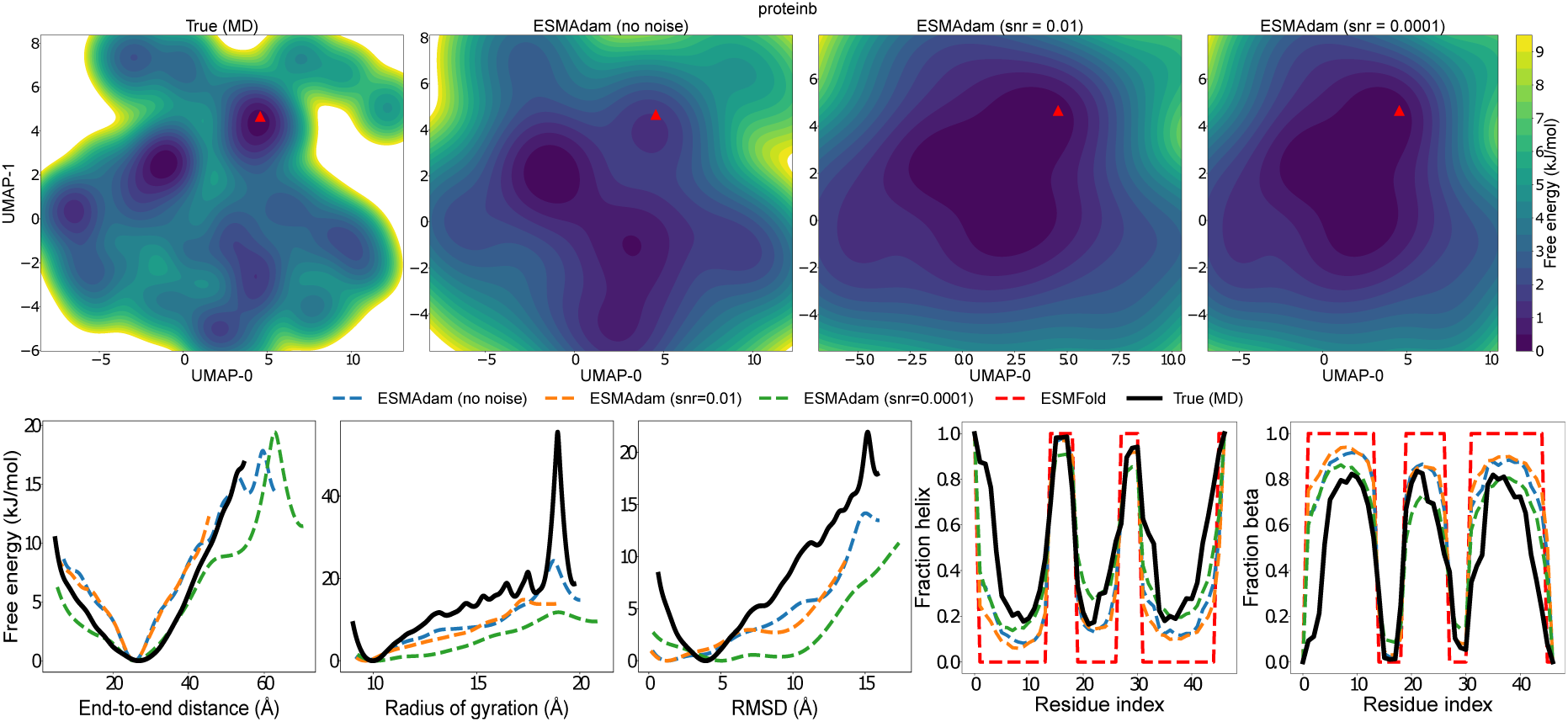
3D conformation ensemble reconstruction from cryo-EM results for ProteinB

**Figure 7:**
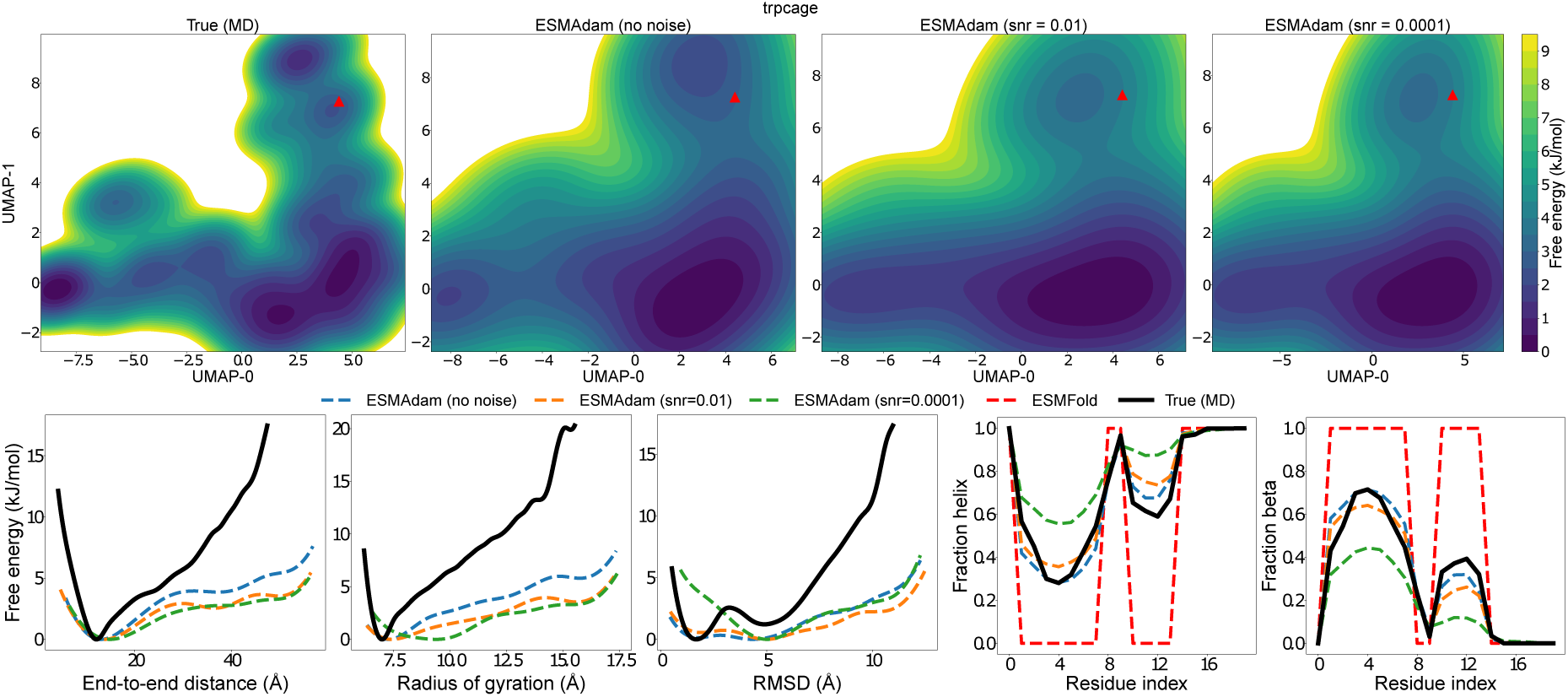
3D conformation ensemble reconstruction from cryo-EM results for TrpCage

**Figure 8:**
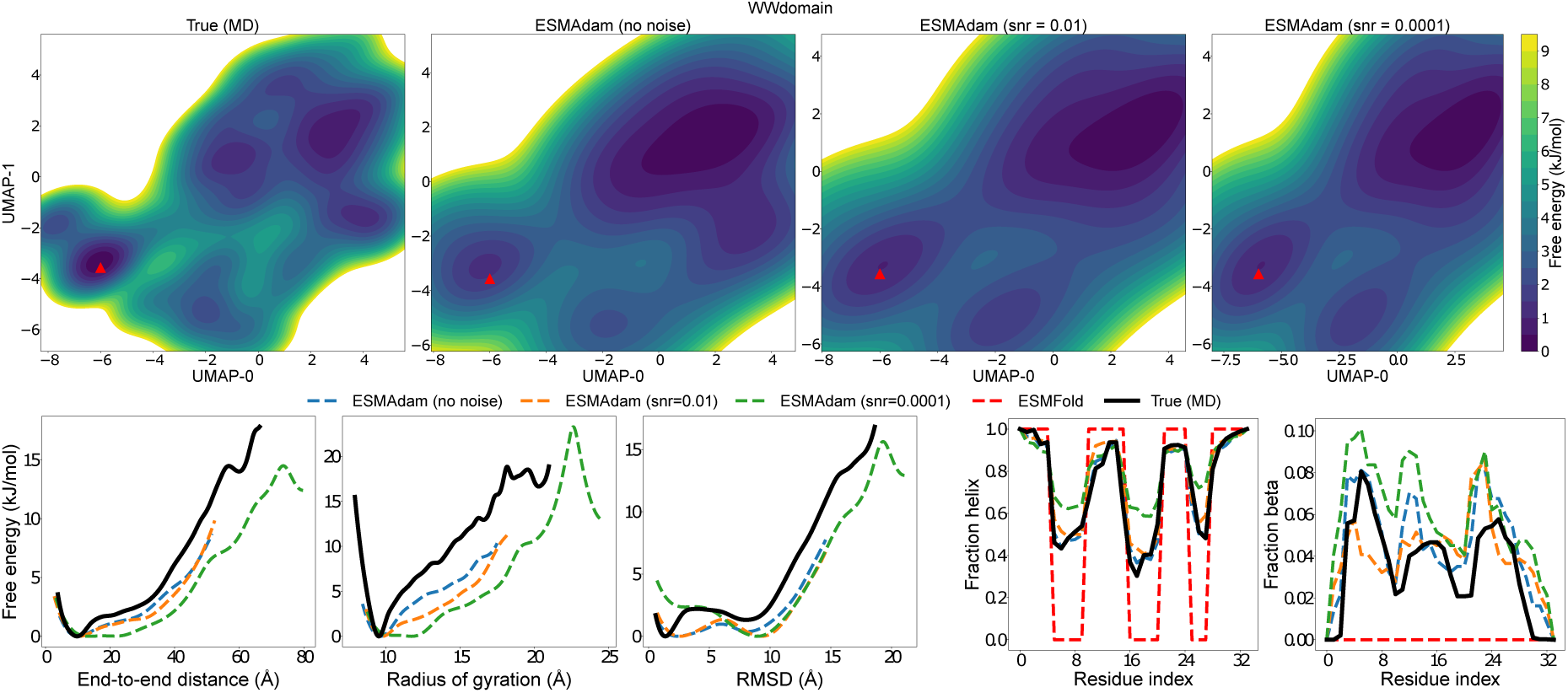
3D conformation ensemble reconstruction from cryo-EM results for WWdomain

