## supporting information for "ESMAdam: a plug-and-play all-purpose protein ensemble generator"

### Benchmark Protein Dataset

We assemble a testing dataset of 12 proteins, from order proteins to fast-folding proteins to intrinsically disordered proteins, as shown in Fig. S1. The ground truth Boltzmann distribution was obtained from long unbiased MD simulations. Simulations of BBA, BBL, WW domain, Homeodomain, Protein B, and TrpCage were conducted in previous work using the CHARMM22* force field and TIP3P water model at the temperature of 350KThe simulations were performed using adaptive sampling strategy. Simulations of drkN, PaaA2, BPTI,gb3, and Ubquitin were conducted by DE Shaw research using the a99SB-disp force field for 30 $\mu s$. The force field parameters were optimized based on experimental measurements to achieve high accuracy in simulation results across various ordered and disordered proteins. The simulation of RS-peptide consists long unbiased simulations of up to 2 milliseconds of simulation time, with 20 unique starting structures, each run with 100 clones of ∼0.9 micro-second each using Amber03ws force fieldand Tip4p/2005 water model. Details of RS-peptide simulation can be referred from previous works. Finally, the simulation of Barnase–barstar protein complex system is conducted by DE Shaw research with Amber ff99SB*-ILDN force field and the TIP3P water model using tempered binding simulations.[1]

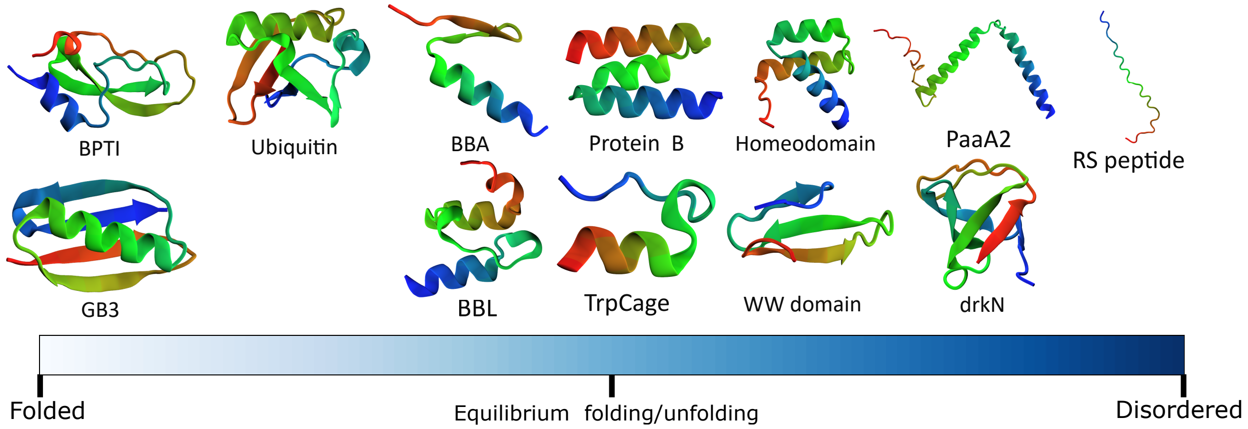


Figure S1: Schematic illustration of benchmark set of proteins used in this work to assess the performance of ESMAdam.

### Full experimental results

**Condition protein conformation ensemble generation** Here we present the full evaluation of the experiment, including the physical feature FES in Fig. S2 and secondary structure percentage per residue in Fig. S3.

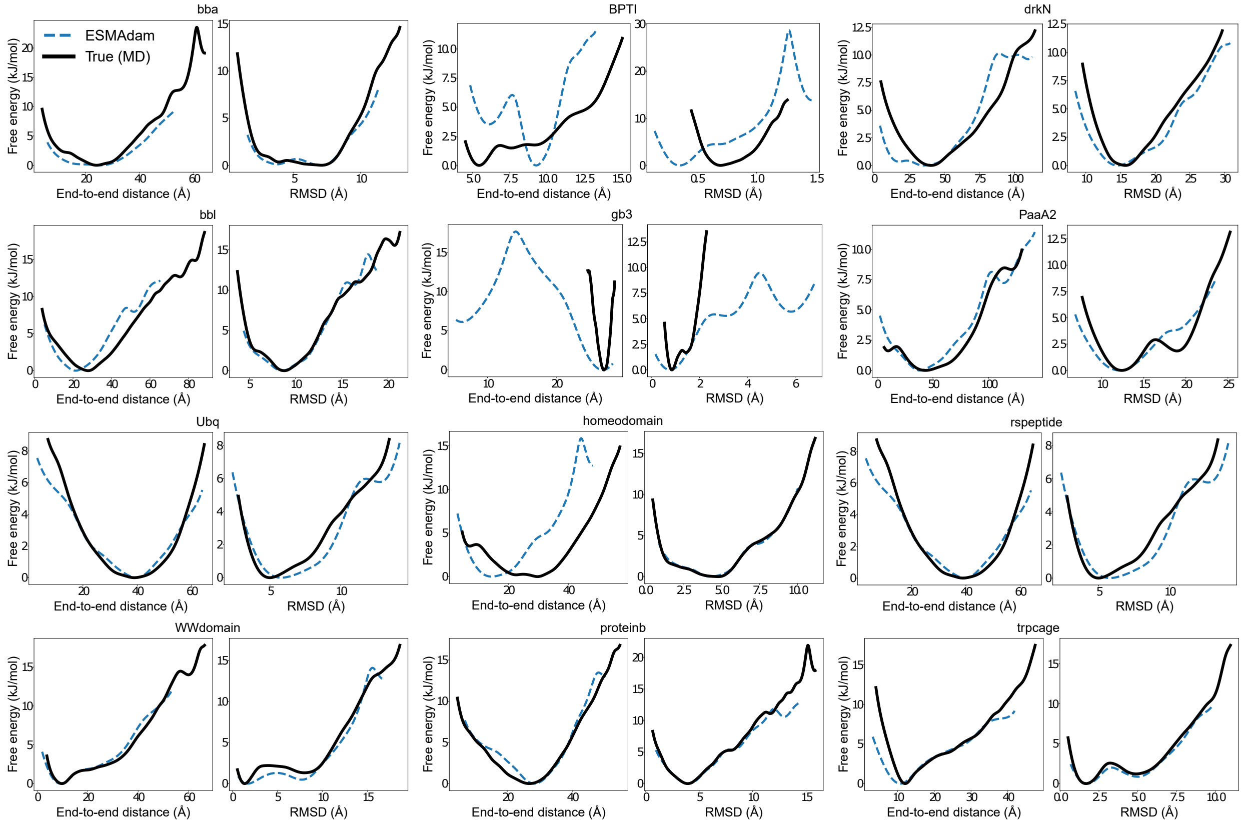


Figure S2: The FES of physical features in all conditional conformation ensemble generation experiments.

**CG-to-all-atom backmapping experiments** Here we present the full evaluation of the ultraCG backmapping experiment, including the UMAP 2D FES in Fig. S4.

**3D structure reconstruction from cryo-EM images** Here we present the additional 3D structure reconstructionn from cryo-EM images experiment for four other proteins: BBL, ProteinB, TrpCage, and WWdomain. The results are shown in Fig. S5, Fig.S6, Fig. S7, and Fig. S8.

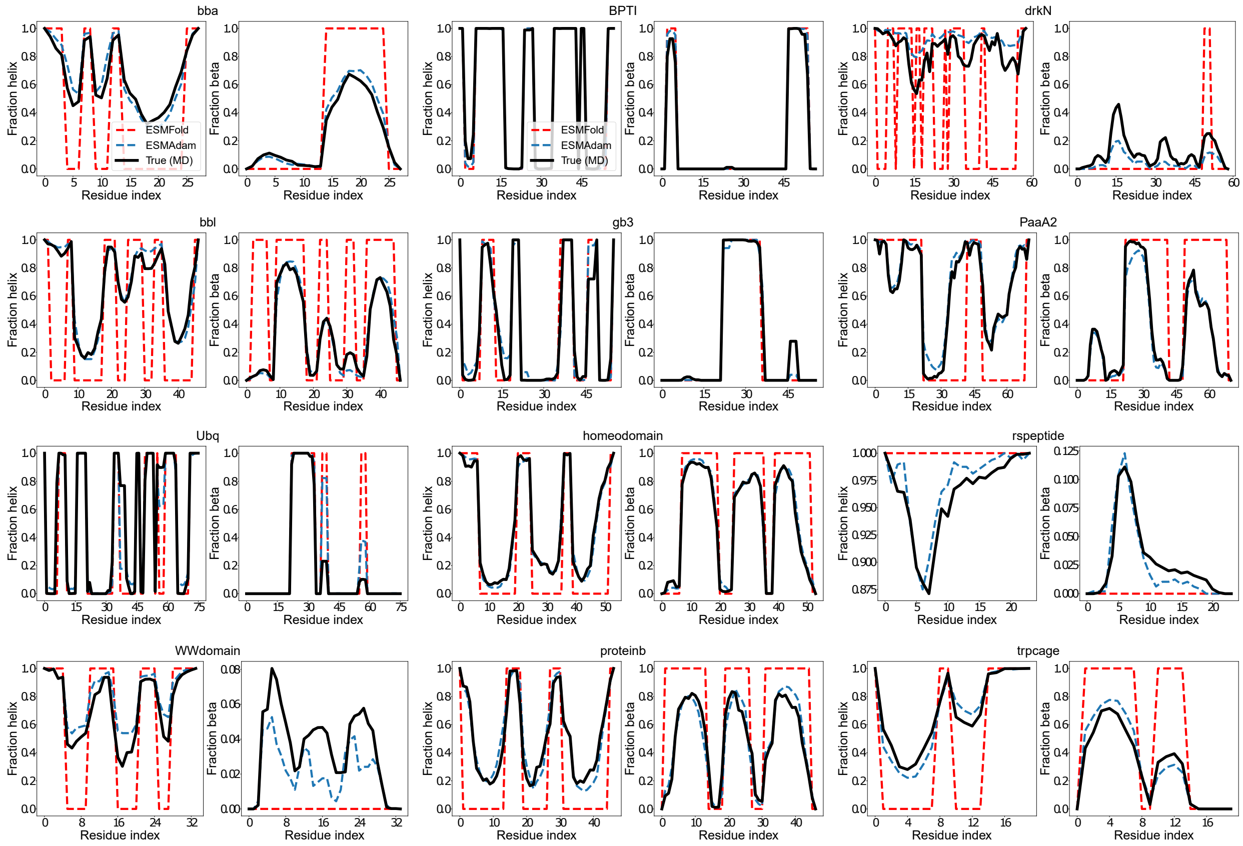


Figure S3: The secondary structure per residue in all conditional conformation ensemble generation experiments.

### Other Methodological Details

**UMAP parameterization** To analyze protein conformational ensembles, we leverage the backbone torsion angles ($\phi$ and $\psi$) as key structural features. These angles are transformed into a high-dimensional feature space by computing their sine and cosine values to ensure a continuous representation. We represent the protein's conformational landscape with this high-dimensional representations. To reduce the dimensionality of this space and facilitate visualization, we apply Uniform Manifold Approximation and Projection (UMAP). UMAP is a non-linear dimensionality reduction technique that preserves the local and global structure of the data while projecting it into a lower-dimensional space. By mapping the high-dimensional torsion angle features to a 2D projection, UMAP enables intuitive exploration of the conformational diversity and clustering of protein states. The UMAP function is pa-

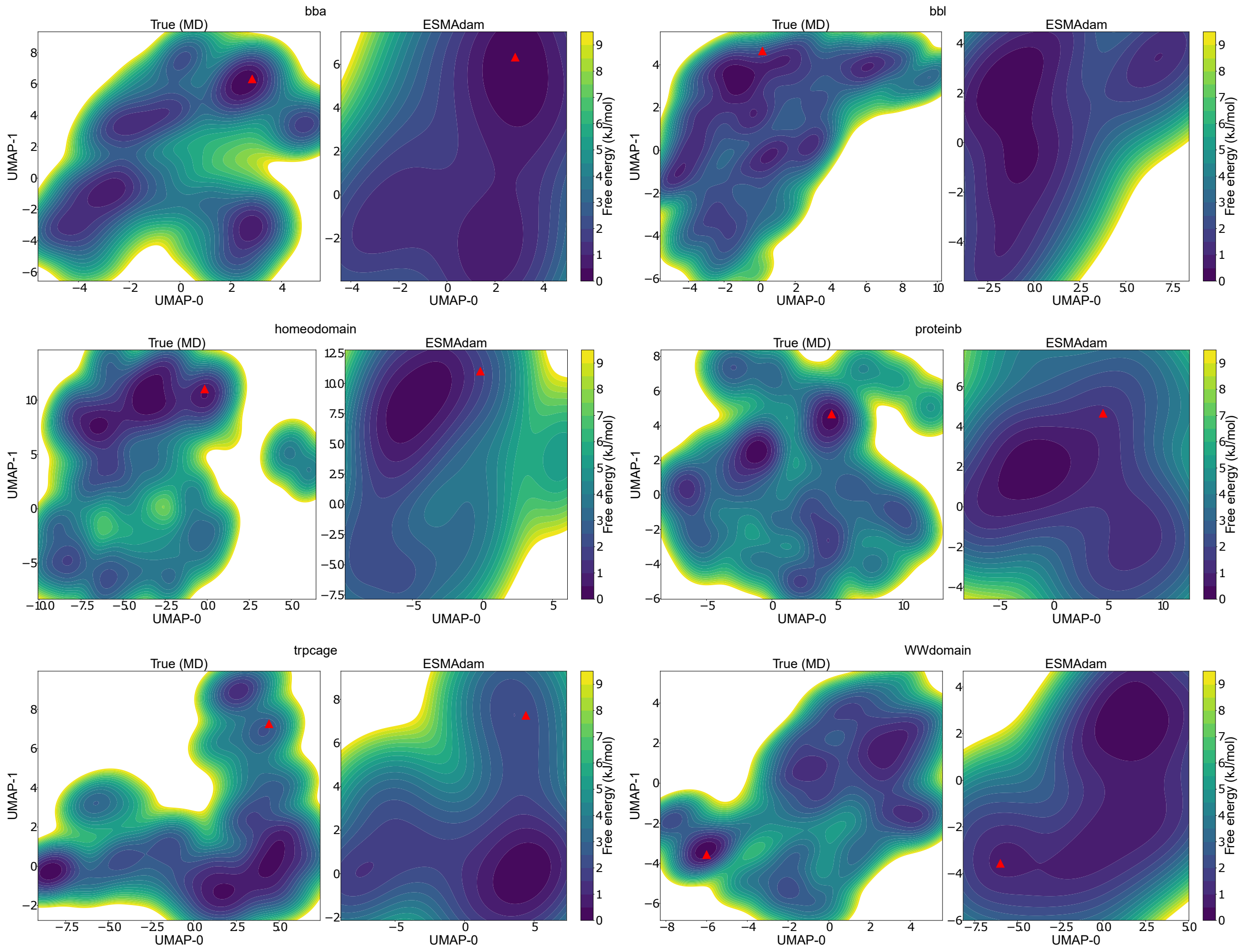


Figure S4: Comparison between the reference MD simulation (left) and ESMAdam (right) in ultraCG backmapping experiment FES across the two dimensional UMAP for each protein.

rameterized with data from the reference MD simulations in all cases.

**Choice of the correction hyperparameter** An important hyperparameter in ESMAdam is the learning rate. A learning rate that is too low can result in prolonged convergence times, while an excessively high learning rate can cause optimization instability. In this study, we report the empirically determined learning rate values that yield the best performance in Table S1. In most experiments, these empirically chosen learning rates enabled convergence within 100 epochs. However, for the 3D structure reconstruction experiments, convergence required up to 300 epochs, reflecting the increased complexity of this task.

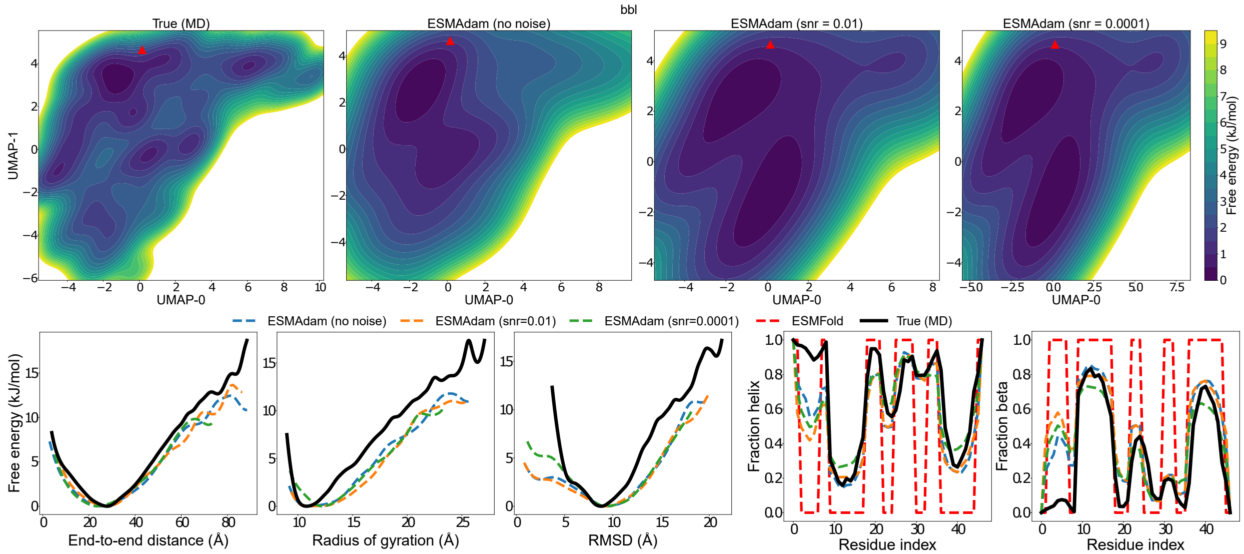


Figure S5: 3D conformation ensemble reconstruction from cryo-EM results for BBL

**Table S1: The correction term hyperparameter for each type of proteins.**

| | Orded | Fast-folding | Disordered |
|---|---|---|---|
| $lr$ | $0.025 - 0.05$ | $0.05 - 0.1$ | $0.1 - 0.25$ |

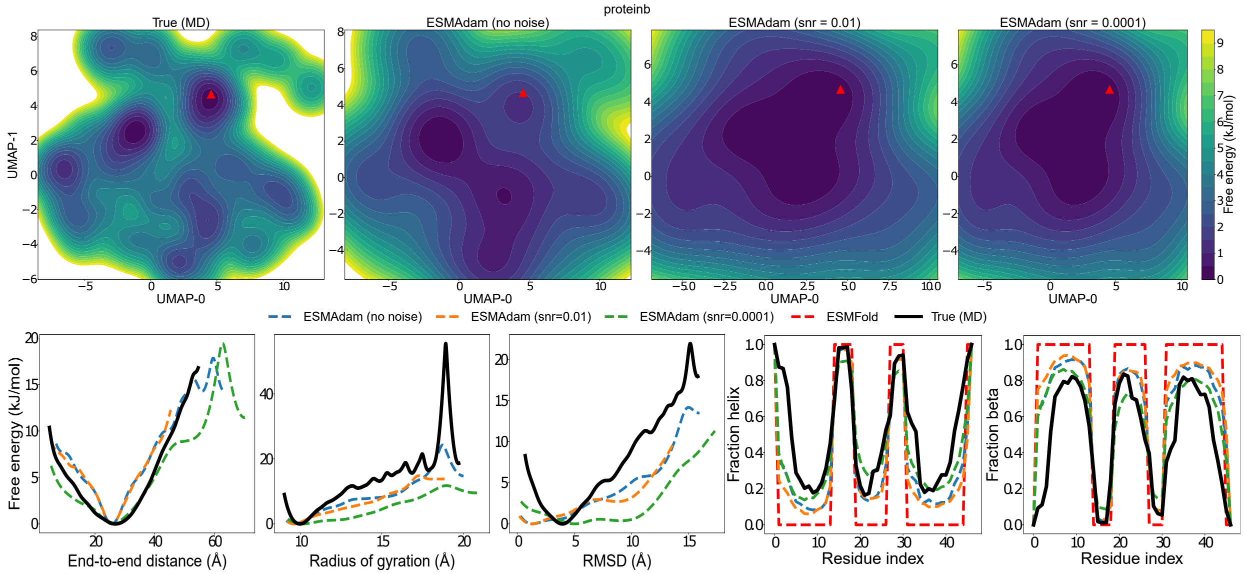


Figure S6: 3D conformation ensemble reconstruction from cryo-EM results for ProteinB

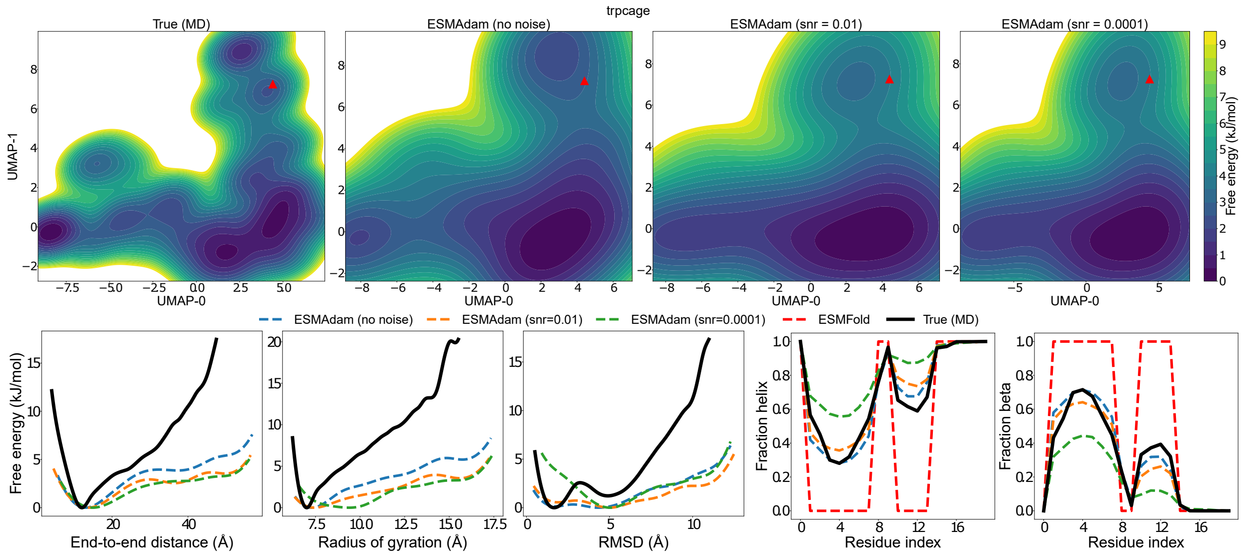


Figure S7: 3D conformation ensemble reconstruction from cryo-EM results for TrpCage

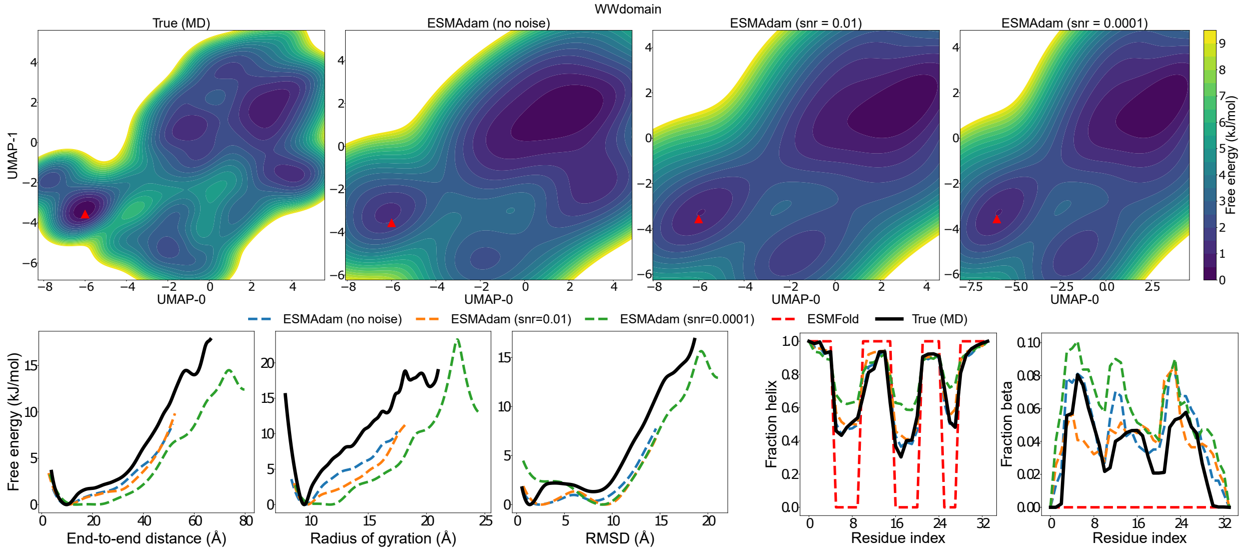


Figure S8: 3D conformation ensemble reconstruction from cryo-EM results for WWdomain
